# Populational Super-Resolution Sparse M/EEG Sources and Connectivity Estimation

**DOI:** 10.1101/346569

**Authors:** E. Gonzalez-Moreira, D. Paz-Linares, E. Martinez-Montes, P. Valdes-Hernandez, Jorge Bosch-Bayard, M.L. Bringas-Vega, P. Valdés-Sosa

## Abstract

In this paper, we describe a novel methodology, BC-VARETA, for estimating the Inverse Solution (sources activity) and its Precision Matrix (connectivity parameters) in the frequency domain representation of Stationary Time Series. The aims of this method are three. First: Joint estimation of Source Activity and Connectivity as a frequency domain linear dynamical system identification approach. Second: Achieve super high resolution in the connectivity estimation through Sparse Hermitian Sources Graphical Model. Third: To be a populational approach, preventing the Inverse Solution and Connectivity statistical analysis across subjects as a postprocessing, by modeling population features of Source Activity and Connectivity. Our claims are supported by a wide simulation framework using realistic head models, realistic Sources Setup, and Inverse Crime effects evaluation. Also, a fair quantitative analysis is performed, based on a diversification of quality measures on which state of the art Inverse Solvers were tested.

## 1 INTRODUCTION

Currently there is a consent at neuroscience community that the brain network connectivity play a crucial role for the understanding of brain functions at behavior and cognition levels by the pattern of communication between brain neuronal regions (*Avena-Koenigsberger, 2018*). In neuroscience field brain networks topology is defined by a group of neural elements (sources) and their interconnections (connectivity) of two kinds: the so called directed or undirected networks (*Salvador, et al., 2005, Estrada, 2012*). Therefore, the brain function characteristics is due to neural groups collective action and mutual interactions of the neural system.

Nevertheless, non-invasive magneto/electroencephalographic (M/EEG) recordings brings an ideal scenario to cover the gap of other “slower” and “indirect” imaging methods (such as fMRI, fNIRS, PET), given its direct association to Neuronal Local Field Potentials and high temporal resolution, which allows the M/EEG to follow at a time-scale of milliseconds and with direct Electrophysiological substrate the Neural events involving human perception and cognition (*Schomer and Lopes da Silva 2011*, *Hämäläinen, et al., 1993*). M/EEG signals are widely used to reconstruct the neural dynamics in both Resting State (RS) and Event Related Potential (ERP). These signals carry the effect of multiple sources within the gray matter, thus M/EEG connectivity analysis constitute a strong asset for noninvasively study brain functional networks (*Schoffelen and Gross, 2009*, *Smit, et al., 2008*).

Unfortunately, M/EEG Sources and Connectivity analysis has proved to be a tough problem. The reason for this is that M/EEG are an smeared projection at the scalp of sources in the brain. Consequently, what we observe at the sensors are signals as products of mixture due to volume conduction and field spreading (*Hassan and Wendling, 2018*). There are many works in the past pursuing the brain functional connectivity analysis at the scalp level by statistics of the sensor’s signal interdependences (*Blinowska, 2011*, *Kaminski and Blinowska, 2014*). In fact, these approaches fail to have a precise matching with anatomical areas, extracting conclusion about brain information processing without an adequate physiological/anatomical basis (*Papadopoulou, et al., 2015*). Therefore, ignoring the influence of volume conduction these methods “hope” that resulting patterns at the scalp would reflect the underlying brain activity, which have been strongly criticized by neuroscience community during the last years (*Haufe, et al., 2013*, *Brunner, et al, 2016, Van de Steen, et al., 2017*).

Moreover, it’s affirmed that the estimation of connectivity at the generator’s level will be improved by Electrophysiological Source Imaging (ESI) methods in both time and frequency domain, accounting from MNE (*Hämäläinen and Ilmoniemi, 1994*), LORETA family (*Pascual, 1999*; *Pascual, 2002; Haufe, 2016*), sparse methods (*Friedman, 2008*), to ENET-SSBL (*Paz-Linares, 2017*). Mostly based on mathematical or penalty models, anatomical constraints of M/EEG generators and biophysical head conductivity models, they are settled upon highly ill-conditioned mathematical framework which is still an issue.

Up to now, the approaches to estimation of cortical connectivity based on ESI methods work in three main stages, **firstly:** to carry out the inverse solution for a single subject without considering connectivity information which usually is the solution of a Bayesian problem with an arbitrary prior connectivity matrix, **secondly:** perform independently the connectivity estimation by statistical analysis of the Sources’ time series (*Sakkalis, 2011*, *Bastos and Schoffelen, 2016*), such as Granger Causality (*Granger, 1969*), Dynamical Causal Models (DCM) (*Penny, 2004*), frequency domain connectivity measures like Coherence (Coh) (*Tucker, et al. 1986; Srinivasan, et al., 2007*; *Guillon, et al., 2017*), Partial Coherence (PCoh) (*Lopes da Silva, et al., 1980*), Directed Coherence (DC) and Partial Directed Coherence (PDC) (*Baccalá and Sameshima, 2001*), **thirdly:** population statistical analysis of the results for sources activity and connectivity features extraction (*Hipp et al., 2012*; *Babiloni et al., 2005*; *Brookes, 2001*).

Nevertheless, these ESI methods with prior connectivity have been developed mainly to estimate activation and not connectivity. Therefore, the connectivity estimation constitutes a postprocessing after inverse solution computation as it is the consequently population features extraction, so, imprecisions in the source localization and source time series reconstruction strongly affect the results about connectivity and thus population features. Additionally, the brain connectivity is a special case of System Identification (SI) problem in which the whole model variables (sources activity and connectivity features of the population) should be estimated to fit the data, those, separated approaches incur into a severe methodological error.

However, the correct solution to this conceptual problem stands for the need of simultaneously estimation of activation and connectivity given the state-space nature of the M/EEG model (*Galka, et al., 2004, Valdes-Sosa, 2004*), and its subsequent extension to populations. Our research is deep motivated by previous work, Variable Resolution Electromagnetic Tomographic Analysis (VARETA) method, well stablished in Inverse Solution by the estimation of the covariance matrix at the sources level (*Valdes-Sosa, 1996*, *Bosch-Bayard, et al., 2001*). Also, given the recent outbreak in the literature on estimation of high dimension covariance matrices and inverse covariance matrices (*Maurya, 2016*, *McGillivray, 2016*, *Ledoit and Wolf 2015*, *Cai, et al., 2016*, *Adegoke, et al., 2018*), we are encouraged to use similar approach. Here we consider and extension of these previous developments to the complex variable case, which arises from the frequency domain representation of the M/EEG forward model of the population, based on the Complex sources Effective Empirical Covariance matrix and the Precision Matrix, the last carrying the functional brain connectivity as a populational statistic.

In this paper, we propose a novel methodology for Inverse Solution (sources activity) and its Precision Matrix (connectivity parameters) for multiple subject’s stationary time series in the frequency domain by means of populational Hermitian graphical lasso prior. This approach attempts for stablishing a new procedure with three important outcomes: A complete dynamical system identification by joint estimation of Sources Activity and Connectivity, Super-Resolution in connectivity estimation through sparse models upon the precision matrix and Modeling the entire population inverse solution and connectivity features. In the next sections of this paper, we provide detailed theoretical demonstrations of the method and its promising results for realistic scenarios like variable source localization, complex connectivity structure, levels of SNR by using pseudo EEG and real data of EEG and MEG.

## 2 METHODOLOGY: BC-VARETA

In what follows we formulate and provide implementation details of a framework that allows for the joint reconstruction of M/EEG Sources and Connectivity features for multiple subjects. This is done by imposing penalties on the frequency domain populational sources variances and covariances matrix achieving great flexibility and precision upon its estimation. This strategy has been stablished by the well-known VARETA, and this methodological refinement attempt for going to the next step, i.e. Variable Resolution Functional Connectivity Analysis. So, we refer to our methodology as Functional Connectivity Variable Resolution Electromagnetic Tomographic Analysis (**BC-VARETA**)

### 2.1 The ECM Method with Population Connectivity Sparse Penalty Model

Here we describe the technical aspects of the inference through the Expectation Conditional Maximization (ECM) algorithm and maximum a posteriori analysis throughout Sources Graphical Model (SGM) on the inverse covariance matrix (*Rubin, 1977*, *Rubin, 1994*, *Hsieh, 2014*).

#### 2.1.1 The frequency domain MEG/EEG Covariance Components Model in populations

The CCM, represents the relationship between the observed variables or data (***V***) and unobserved variables or parameters (***J***) as a linear model, in a similar fashion to the definition of the MEG/EEG forward model, see equation [1.1]:

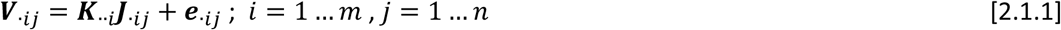

The data (***V***)*_p_*_×*m*×*n*_, parameters (***J***)*_q_*_×*m*×*n*_ and noise (***e***)*_p_*_×*m*×*n*_ are defined as complex number Tensors, from the frequency domain representation (at a single frequency bin) of the M/EEG recordings, sources’ PCD and M/EEG sensors’ noise, respectively, where *p* is the number of sensors, *q* is the number of generators, *m* is the number of subjects under analysis and *n* is the number of time windows in which the Fourier coefficients are computed. Furthermore, the source to sensors design Tensor (***K***)*_p_*_×*q*×*m*_ is obtained from the discretization of the specific head model Lead Field, in a common generators and sensors space across subjects. For the complex noise vectors ***e***_·*ij*_ it is assumed that constitute independent random vectors with Circularly Symmetric Complex Multivariate Normal probability density function (pdf) (*Marzetta, 1995*):

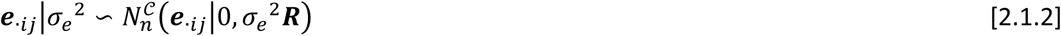

Where nuisance scale parameter *σ_e_*^2^ is an unknown positive scale hyperparameter and (***R***)*_p_*_×*p*_ is a known positive definite and hermitic matrix. Under these assumptions the likelihood or the observed variables *pdf* will be the following:

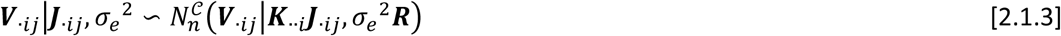

Therefore, the M/EEG Data Empirical Covariance Tensor is defined:

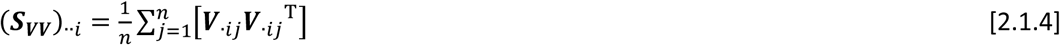

The complex sources’ PCD vectors ***J***_·*ij*_ are modeled as independent random variables with Circularly Symmetric complex multivariate Normal *pdf*, where the covariance or cross-spectral matrix (***Σ_JJ_***) *_q_*_×_ *_q_* is defined as an unknown positive semidefinite and hermitic matrix of hyperparameters to be estimated:

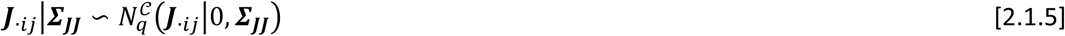

A prior *pdf* over the hyperparameters (*σ_e_*^2^) and (***Σ_JJ_***) could be considered. In this sense it is important to notice that by using prior *pdf* on the cross-spectral matrix (***Σ_JJ_***) we are regularizing the real and imaginary part. For simplicity, all hyperparameters including (*σ_e_*^2^) and (***Σ_JJ_***) are summarized within a unique variable (**Ω**), thus the whole hyperparameters set prior *pdf* can be denoted as *p*(**Ω**). The data and parameters {***V****_·ij_*,***J****_·ij_*}; *i* = 1 … *m*, *_j_* = 1 …*n*, can be rearranged into what is defined as complete data (***V***,***J***), so its joint *pdf* can be referred as complete data likelihood *L_c_*(**Ω**) = *p*(***V***,***J***|**Ω**). The CCM admits a hierarchical representation by means of a Bayesian Network (Figure 1), where the estimation of the unobserved data or parameters (***J***) constitute the first level of inference followed by the estimation of the hyperparameters (**Ω**), or second level of inference.

**Figure 1.**
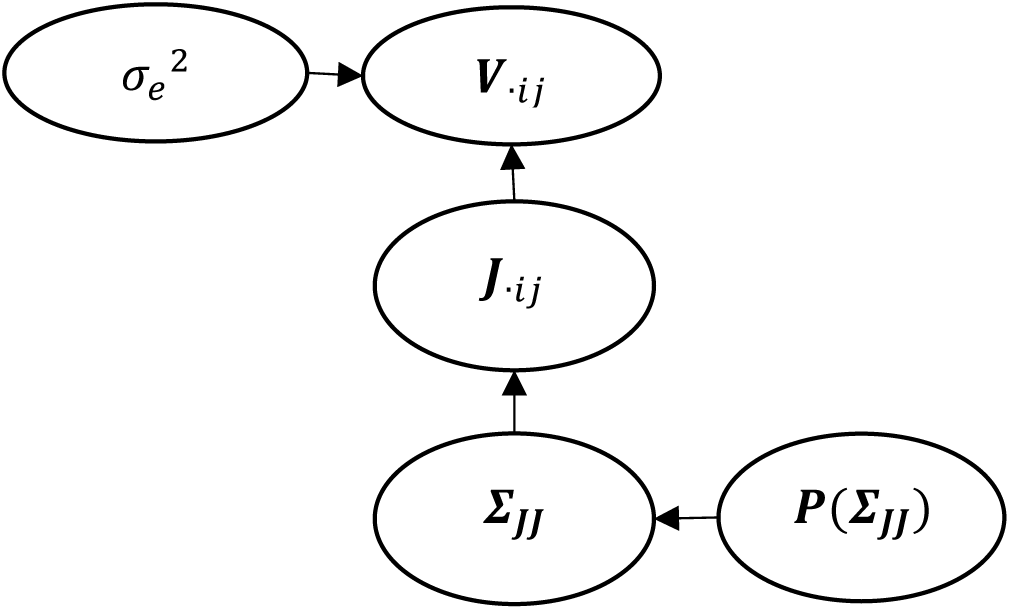
Bayesian network of the covariance Components Model with covariance matrix prior

#### 2.1.2 Population Expectation Step

A Maximum a Posteriori (MAP) analysis of the hyperparameters can be performed by applying the expectation over the unobserved variables of the hyperparameters’ log posterior *pdf* (*McLachlan, 2007*). Using the Bayes rule the hyperparameters’ posterior *pdf* can be expressed as proportional to the product of the Complete Data Likelihood and the hyperparameters’ prior *pdf*:

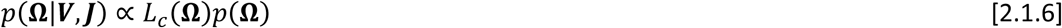

The expectation operation on [2.1.6] can be written in short by formulas [2.1.7] and [2.1.8] below, where (**Ω**^(*k*)^) represent fixed values of the hyperparameters, providing the hyperparameters’ iterated posterior density function:

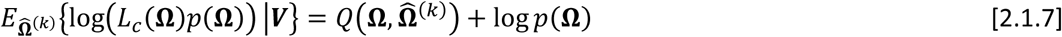

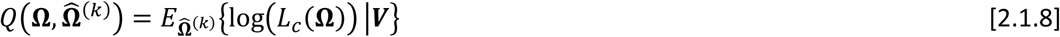

Under the model assumptions of formulas [2.1.3], [2.1.5] and with a fixed value of the hyperparameters (**Ω**^(*k*)^) it is straightforward to show that the unobserved data {***J****_·ij_*}; *i* = 1 …*m*, *j* = 1 …*n* posterior *pdf* is a multivariate Normal distribution with individual subjects posterior mean 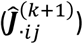 and Covariance matrix 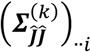 (*see Appendix B1*):

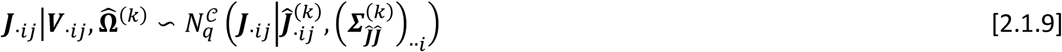

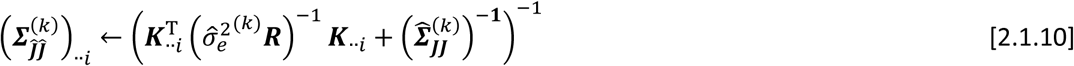

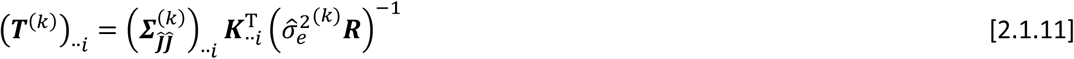

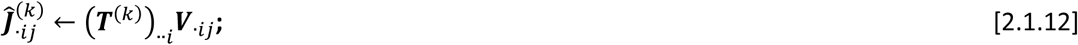

Where the slices of the Tensor ***T*** constitute the denominated data to sources’ *Transference Operator*^1^ for individual subjects.

After some algebraic transformation of the Expectation in equation [2.1.8] by using equations [2.1.9], [2.1.10], [2.1.11] and [2.1.12] (*see Appendix B2*), we can reach to a closed expression of the function 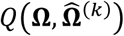:

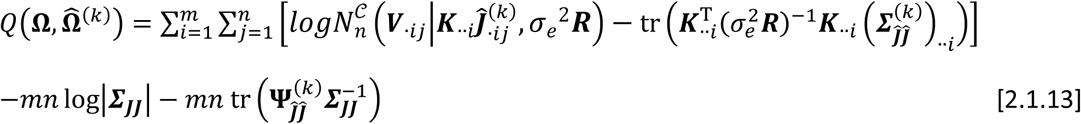

The matrix 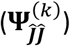, hereinafter we will call the Effective Sources Empirical Covariance (ESEC), since it carries the information about sources correlations that will effectively influence the estimator of (***Σ_JJ_***) in the maximization step, is defined as the sum of two components, the subjects mean of the sources (parameters) posterior covariance 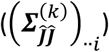 and the Sources Empirical Covariance 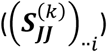:

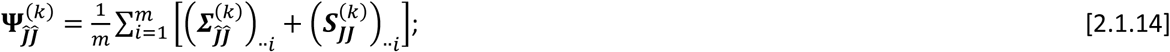

Where the Source Covariance 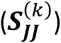 is defined:

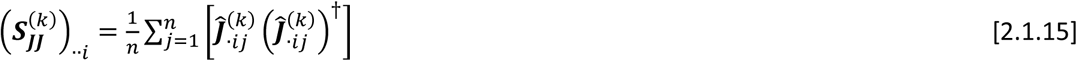

In virtue of [2.1.12] the ESEC can be expressed as a function of the data empirical covariance (***S_VV_***):

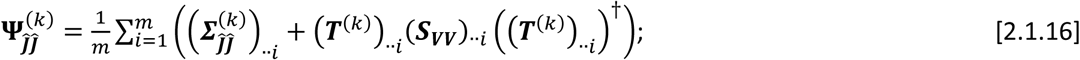

#### 2.1.3 Population Maximization Step

The estimator of the hyperparameters 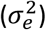 can be obtained by equating to zero the corresponding derivative of [2.1.13]. If for a better numerical control, we consider using a Jeffreys Improper Prior *pdf* (*Jeffreys, 1946*) with shape 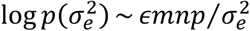, the following estimator arises (*see Appendix B3*):

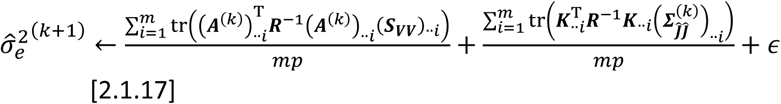

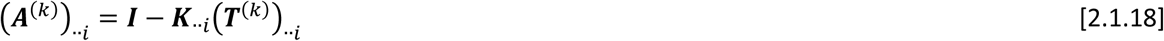

Without losing generality the prior *pdf* of (***Σ_JJ_***) can be expressed as an exponential of a penalty function *P*(***Σ_JJ_***, *λ*), so that log*p*(***Σ_JJ_***) ~ – *mP*(***Σ_JJ_***, *λ*), where the coefficient *m* has been set for convenience and *λ* represents a vector of regularization parameters or tuning hyperparameters of the covariance matrix prior *pdf*. After substituting log*p*(***Σ_JJ_***) in [2.1.7], eliminating the factor *m* and changing the sign, the maximization step over (***Σ_JJ_***) turns into the minimization of the auxiliary target function:

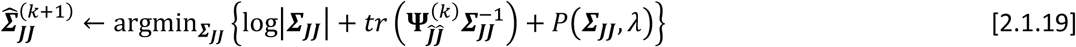

The estimation of the regularization parameters will be done iteratively by maximizing the likelihood through direct differentiation of the terms dependent on (*λ*) [2.1.6] and setting a prior *p*(*λ*) that guarantees a close form solution and the existence of a single local maxima at every iteration (*see Appendix D1*):

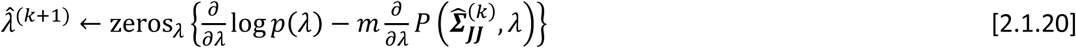

The hyperparameters’ iterated posterior distribution provides a target function for the estimation of the covariance matrix that is independent on the rest of the current hyperparameters estimators. Importantly, it can be seen that [2.1.20] is dependent through ESEC from the hyperparameters obtained in the previous maximization step and from the parameters and its posterior covariance estimates given in the current expectation stage. The sources estimation or first level of inference will influence downwards into the second level of inference, estimation of the covariance matrix, through the iterated ESEC, and the estimation of the covariance matrix and its properties will affect upwards into the model inference, sources estimation. The estimation of the covariance matrix becomes a separated problem itself that we describe in the next section.

### 2.2 The Penalized Population Inverse Covariance model

Here we describe a methodology for the second level of inference or Sources’ Graphical Model (SGM) estimation, similar to the typical Graphical Model (*Whittaker, 2009*), where the ESEC performs analogously to the empirical covariance in this context but in the sources’ level. We reformulate the target function as a function of the Inverse covariance and incorporate a Penalized Inverse Covariance model (PIC) as a prior pdf in the BC-VARETA Maximization step. The PIC model when combined with a strategy for SGM estimation based on Proximal Newton-type Methods (*Schmidt, 2010*) lead us to an iterative scheme where the computation of the descend direction is tackled into the Penalized Least Squares (PLS) framework.

#### 2.2.1 Population Sources’ Graphical Model with Penalized Inverse Covariance

The penalty function *P* in equation [2.1.19] is redefined as dependent of the inverse covariance (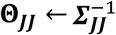):

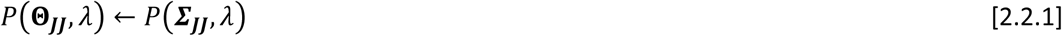

When the target function is redefined over the Inverse covariance matrix by changing variable (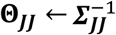) within the whole expression [2.1.19] we obtain an equivalent minimization of Graphical Model type target function instead:

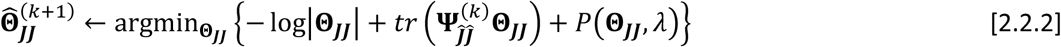

In the context of PIC, a single penalty models leading to different independent estimation schemes reported in literature can be regarded. For example, if we consider the special case of a non-informative prior over the inverse covariance matrix (or constant penalty function *P*(**Θ*_JJ_***) = *cte*), which is equivalent to the implementation of EM by (*McLachlan, 2012*), then the equation [2.2.2] has a unique and closed form solution with the ESEC matrix (*see Appendix B3*).

The Laplace prior with matrix L1 norm exponent or real LASSO model (*P*(**Θ*_JJ_***) =*λ*∥**Θ*_JJ_***∥_1,**Λ**_), where **Λ** is a known positive weights’ matrix representing inverse covariance elementwise precisions, has been used to pursue sparse estimation of the inverse covariance. This model, originally identified within literature as the Graphical LASSO (*Friedman, 2008*, *Mazumder, 2012*), has been tackled by means of several algorithms. They attempt to optimize the functional [2.2.2] under LASSO model by solving the equation that arises from its direct differentiation^2^:

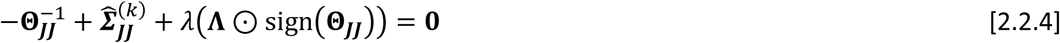

For an implementation of PLS methodology we consider the L1 norm model. It leads to a solution of this Graphical Model optimization problem by LASSO regression (*Tibshirani, 1996*) (*see Appendix C1*). Furthermore, in *Appendix D* we provide with the detailed technical insights into BC-VARETA implementation and its pseudocode which could be summarized in eight main steps:

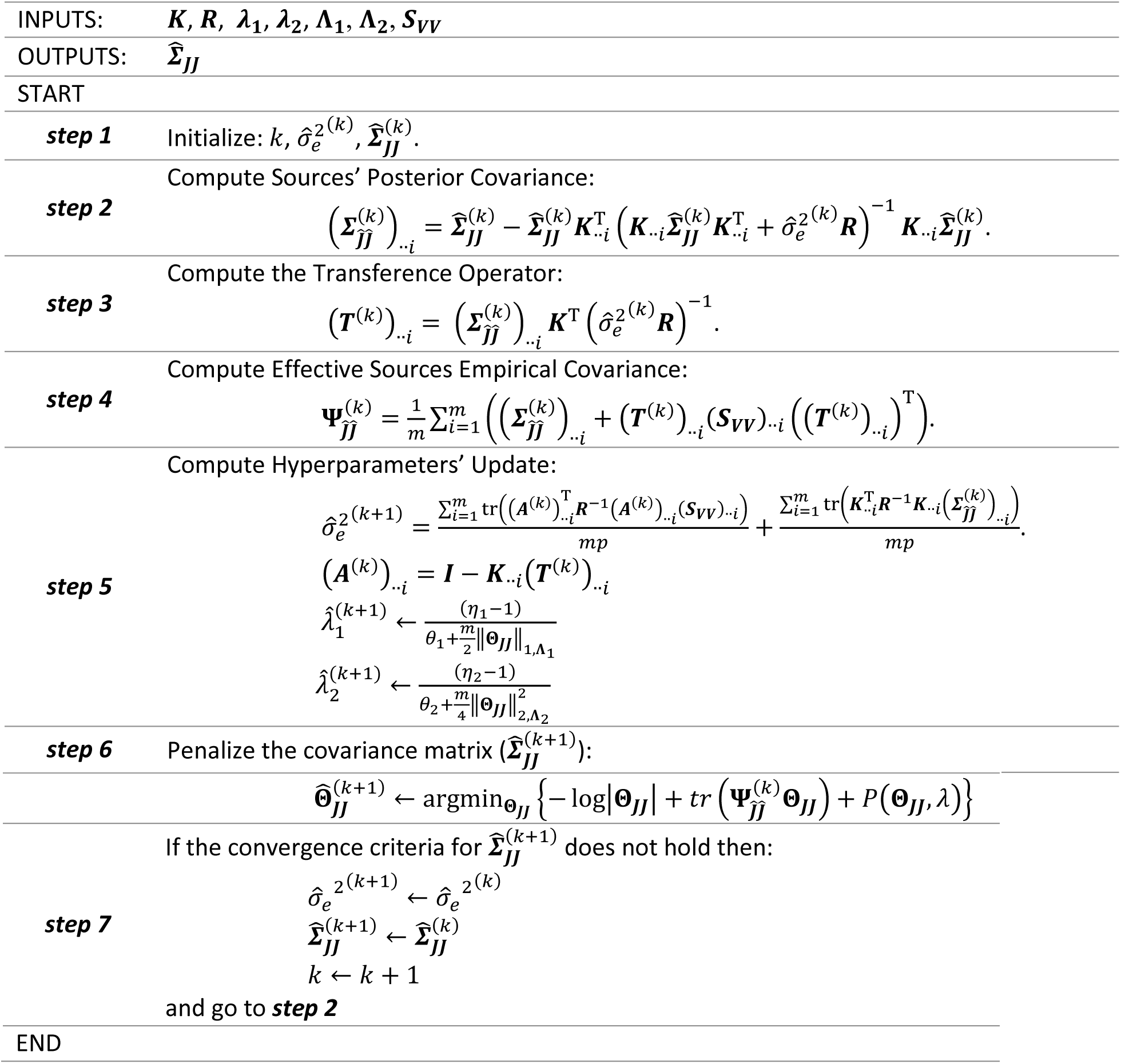

## 3 RESULTS

### 3.1 Simulation study

Traditional inverse solution methods attempt to estimate the localization of the active dipoles and functional connectivity computations is done afterwards. Differently, BC-VARETA method aims to simultaneously estimate, through the effective sources empirical covariance, the activity at sources level and the functional connectivity between the active generators. The main concept behind BC-VARETA method lies over the principle that the activity at the brain bias the brain connectivity and vice versa, therefore the estimation of the inverse solution without taking in account the connectivity could lead to a wrong statistic of the inverse solution. Furthermore, BC-VARETA is a method developed to for variable resolution sources estimation of spatially distributed Neural Activity along the cortical surface, so a fair validation should cover a variety of scenarios of multiple sources with different spatial extensions, different distances between sources centroids and also different connectivity modes.

To analyze the performance of the BC-VARETA method we created a simulation framework representing a realistic scenario of spatially distributed sources (patches) with variable extensions and a dense connectivity pattern within the patches, but attaining for different degrees of sparsity in the connections between them (Figure 2). The generators were defined over a cortical surface of 2004 equally distributed electric dipoles. From the ICBM152 model the dipole position was obtained by down-sampling the 74924 vertices in MNI space coordinates (*Haufe, 2018*).

**Figure 2.**
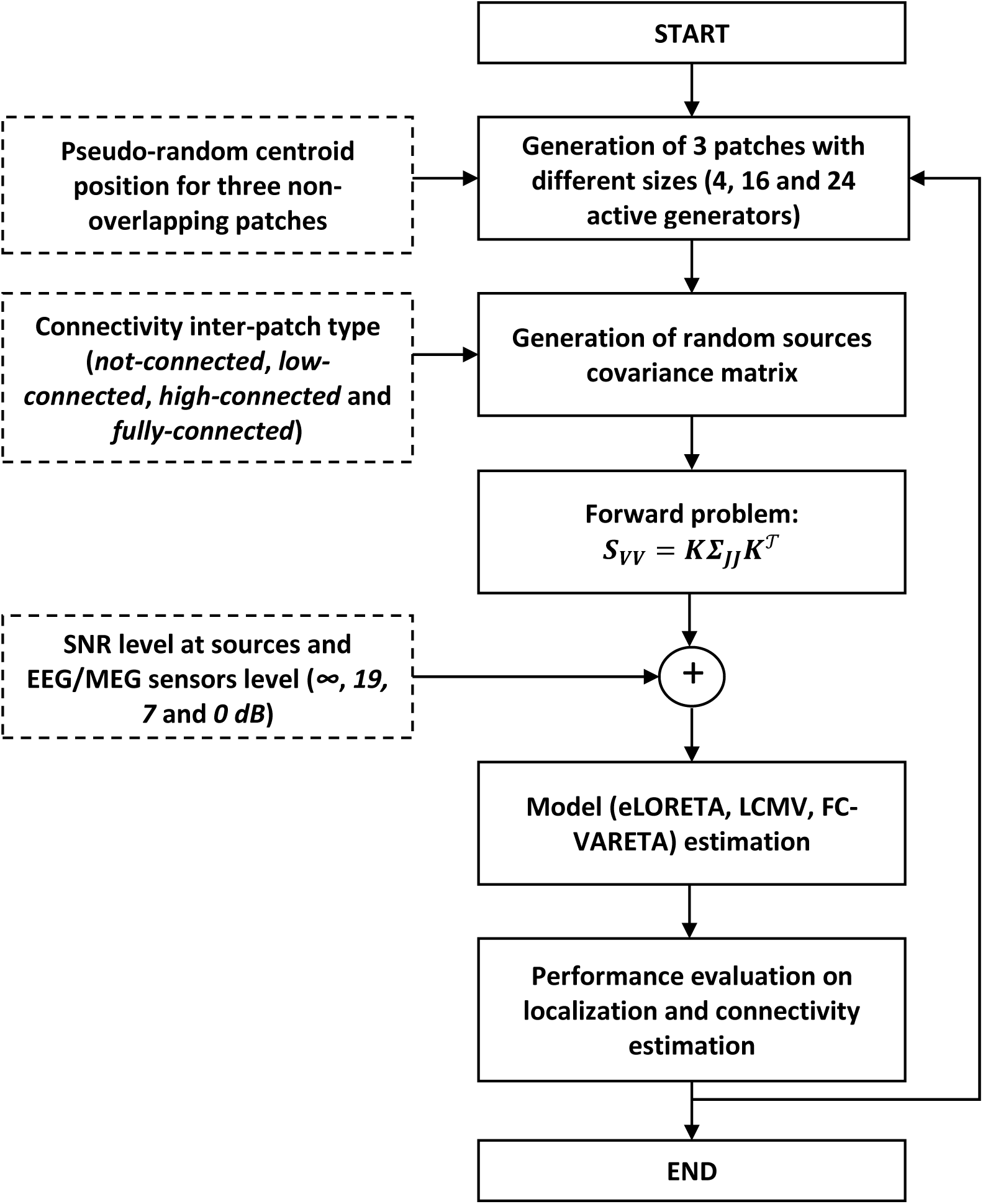
Block diagram of the simulation framework performance evaluation of the connectivity estimation method. The data empirical covariance matrix, ***S_VV_***, defines sensor’s activity (elements on the main diagonal) and covariances (elements out of main diagonal) at the scalp level, meanwhile the covariance matrix, ***Σ_JJ_***, defines the activity (elements on the main diagonal) and connectivity (elements out of main diagonal) at the generator’s level.

Three patches were generated with different sizes (4, 16 and 24 active generators) over the brain cortical surface. The patch centroids were randomly generated into four Regions of Interest: ROI 1, ROI 2, ROI 3 and ROI 4 (Figure 3). However, the localization of the patch centroids was selected according to two main criteria: *close-distance* and *far-distance*. The *close-distance* was defined so that the three patches belonged to the same ROI and without overlapping, which guaranteeing a maximum distance between patches lower than 5 cm and the *far-distance* was defined so that each patch belong to different ROI’s, which guarantee for the distance between patches to be higher than 8 cm.

**Figure 3.**
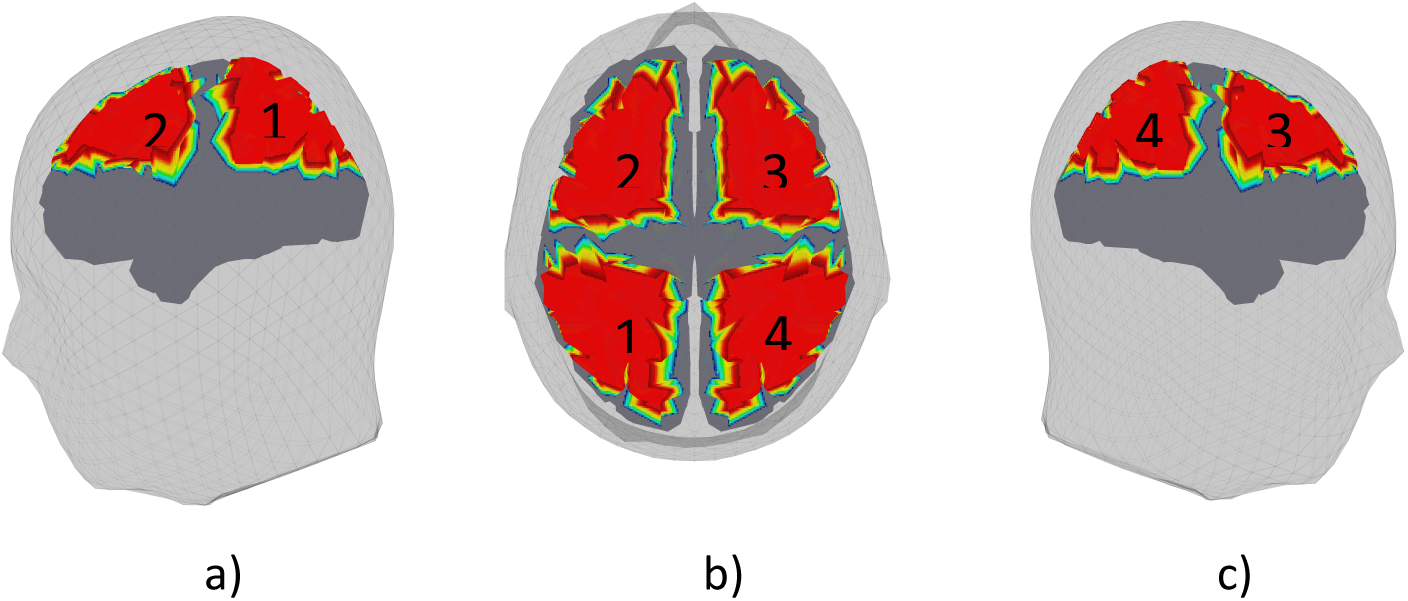
Four ROIs over the ICBM152 brain: a) left hemisphere view, b) axial view and c) right hemisphere view.

Furthermore, the connectivity between the active generators of each patch were defined by four connection modes (Figure 4):

- *not-connected*: all three patches are unconnected between them.
- *low-connected*: first patch and third patch are connected between them, but the second patch is unconnected with first patch and third patch.
- *high-connected*: first patch is connected to second patch and to third patch but there is no connection between second patch and third patch.
- *fully-connected*: all three patches were connected between them.

**Figure 4.**
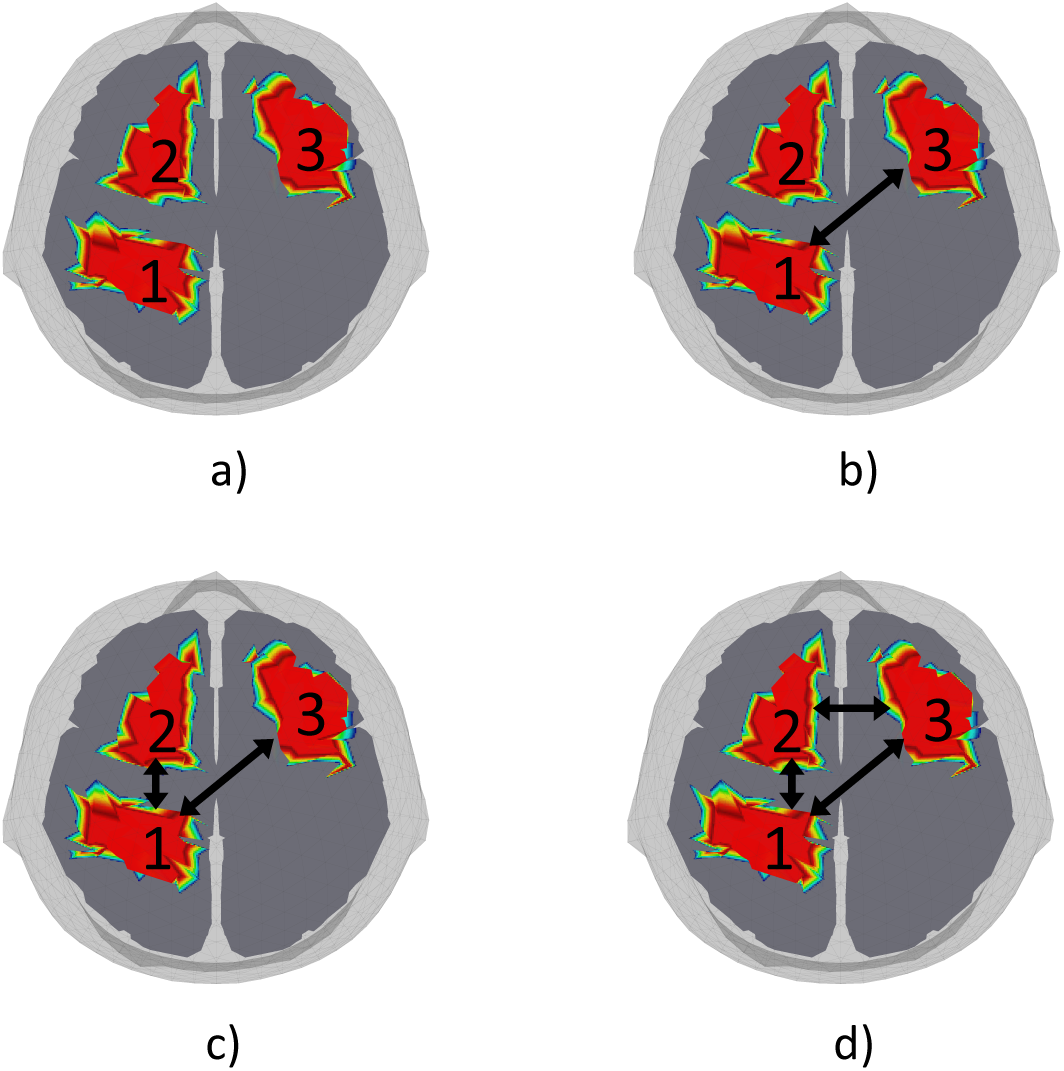
Connection modes between cortical patches: a) not-connected, b) low-connected, c) high-connected and d) fully-connected.

For each of the patches configurations and connectivity modes, the sources covariance matrix (***Σ_JJ_***) was generated with random complex numbers (Figure 5) and it was projected onto 108 EEG electrodes (***S_VV_***) defined by the New York Head model (*Haufe, 2015*, *Huang and Haufe, 2015*) following next equation:

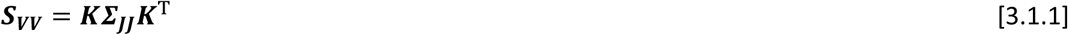

**Figure 5.**
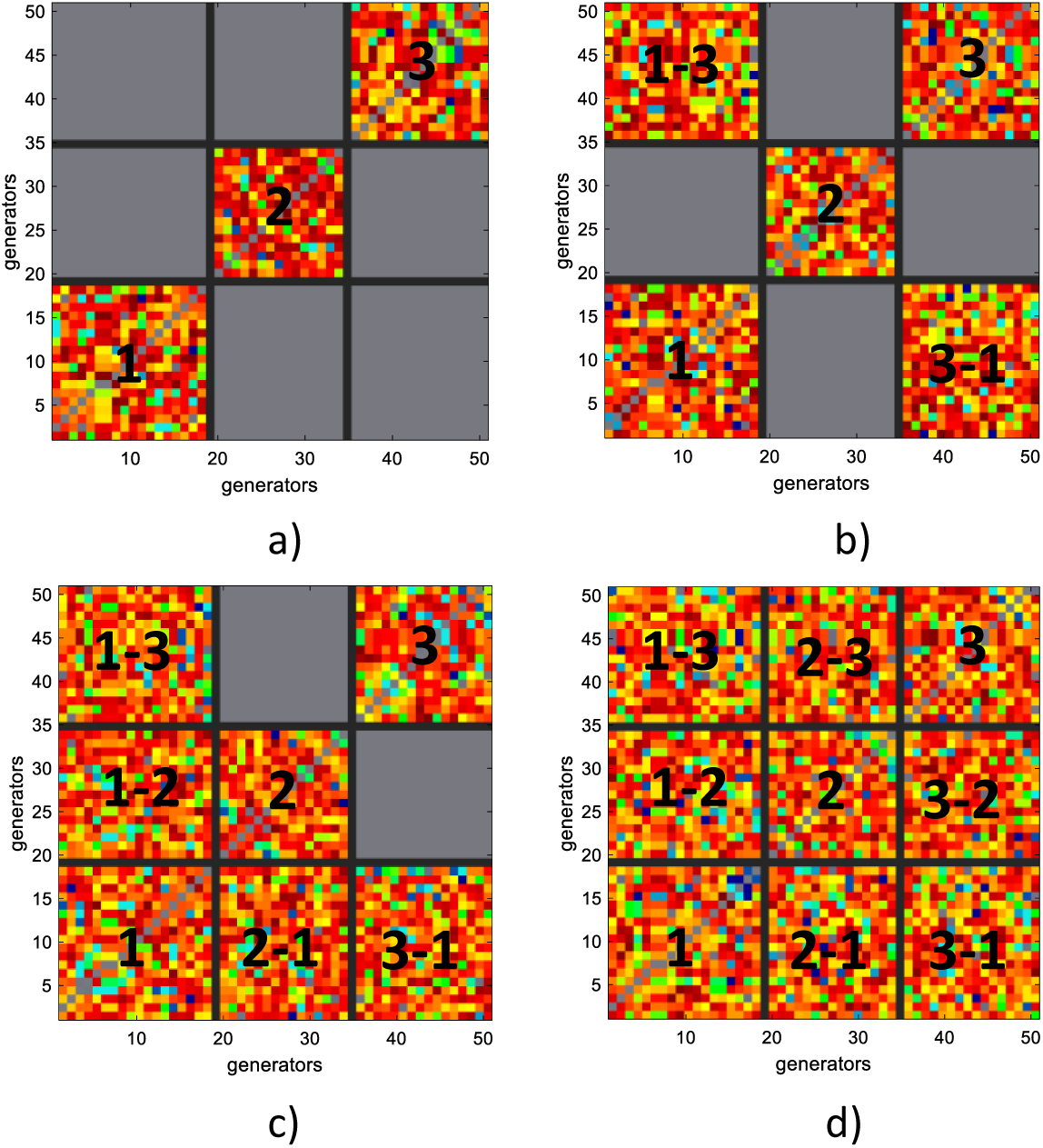
Examples of sources covariance matrix for four connection type between three patches: a) not-connected, b) low-connected, c) high-connected and d) fully-connected.

Furthermore, four different *levels of noise* (∞, 19, 7 and 0 dB) were added to the data covariance matrix, ***S_VV_***. First, the simulation was corrupted with biological noise in 500 generators, following and autoregressive model filtered to create a pink noise, that was projected to the scalp sensors with the M/EEG forward model. Secondly, all sensors (108 electrodes) were contaminated with noise (white noise), mimicking a real situation of an M/EEG recording system.

### 3.2 Simulate data analysis

#### 3.2.1 Localization performance

To evaluate the BC-VARETA performance of variable resolution source retrieval we generate 400 configuration of patches localization (*close distance* and *far distance*). For comparison purpose we use well stablished methods for inverse solution: Linearly Constrained Minimum Variance (LCVM) and Exact Low-Resolution Tomography (eLORETA). LCMV approach is a spatial filtering method that relate the underlying neural activity to the distribution of potential measured at the surface assuming stationary source distribution (*Van Veen et al., 1997*). On the other hand, eLORETA is one of the member of the LORETA family with zero error localization for one active dipole (*Pascual, 2007*).

In Figure 6 we show two examples of solutions for active generator localization over the cortical surface, obtained by eLORETA, LCMV and BC-VARETA. There eLORETA solution reach a completed estimation of the simulated patches but as a typical smoothed estimation it spreads over the cortical surface recovering the larger number of wrong active dipoles. LCMV scenario is more favorable since it is able to recover the correct localization of the three patches, but its solution is not sparse enough to identify the three patches separately. Nevertheless, the estimation reached by BC-VARETA is the sparsest one, and qualitative different to the previous solutions, identifying more accurately the three patches extensions. This simple qualitative analysis constitutes a partial demonstration of our aims: Variable Resolution Estimation, that shall be demonstrated with proper quantitative analysis in what follows. For a complete analysis we evaluate the estimated sources covariance matrix (***S***) by direct measures of its difference with the simulated source covariance matrix (***Σ_JJ_***) (*Ground Truth*). The quantitative evaluation of each result was performed by means of five measures: Sensitivity (TPR), Specificity (TNR), Area Under ROC Curve (AUC), F1 score (F1S), Dipole Localization Error (DLE).

**Figure 6.**
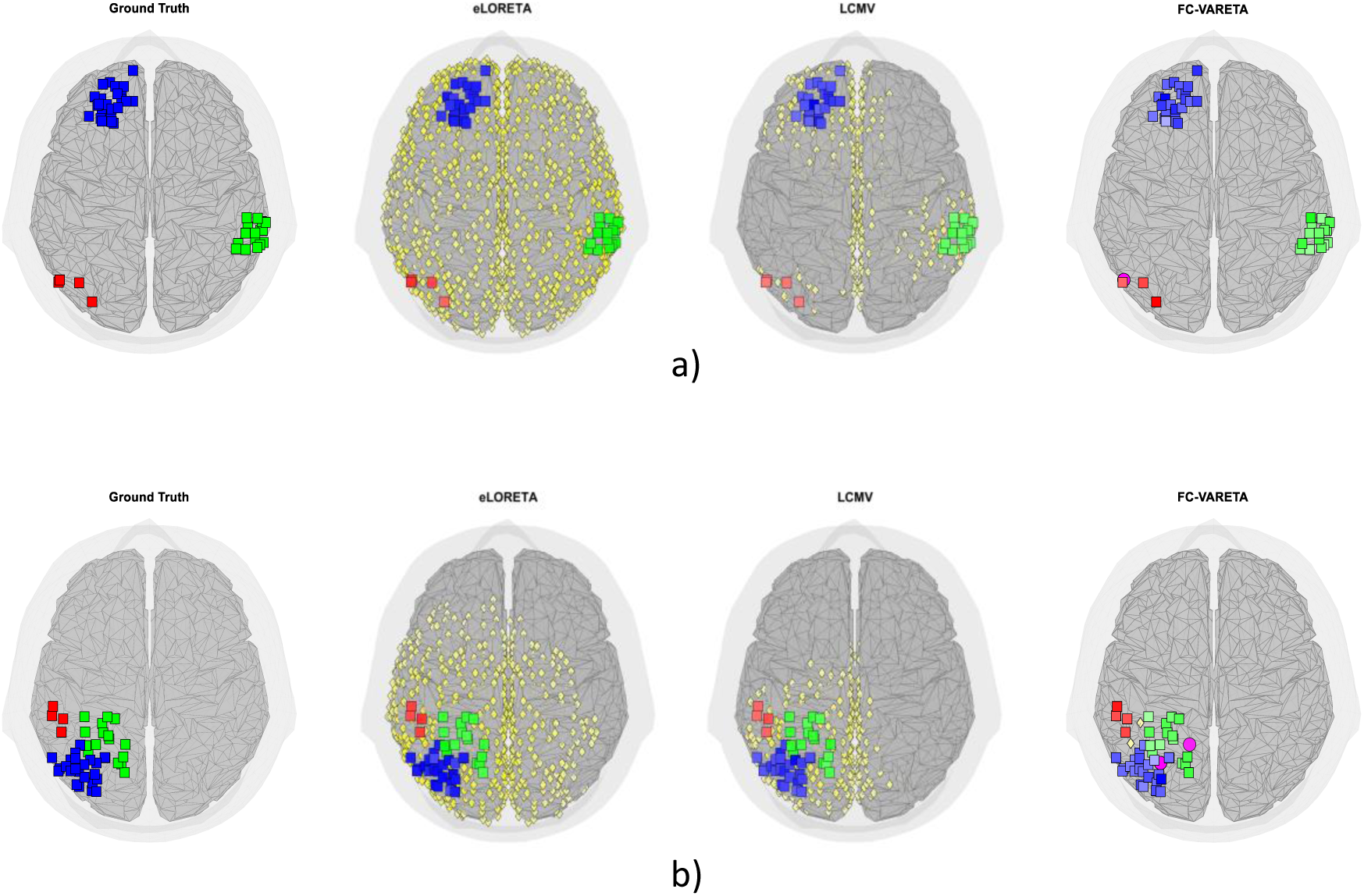
Two simulation configurations of the three patches with different size (4, 8, 24 active generators) for the two spatial distribution criteria: a) close-distance and b) far-distance. The Ground Truth and it localization estimation based on eLORETA, LCMV and BC-VARETA is shown from left to right. The square represents the True Positive estimation (red color for patch 1, green color for patch 2 and blue color for patch 3), the yellow diamond represents the False Positive estimation and the magenta circle represents the False Negative estimation. All the solutions were normalized and values lower than 0.01 (1%) were set to zero.

The results of eLORETA, LCMV and BC-VARETA localization performance by using ROC analysis and DLE estimation are reported in Table 1. The worst results are obtained by eLORETA method since the sparsity level of it inverse solution is lowest, therefore the method doesn’t have the resolution to estimate patches at different size even when the *far-distance* criteria hold. One interesting point to be noted here is that the DLE values for eLORETA solutions are not zero as it is claimed in previous publications, this is due to the fact that eLORETA even when it is able to achieve exact localization for a single simulated dipole, fails to recover multiple sources in a wide range of conditions as shown in Figure 6. In general, for several active dipoles the eLORETA source estimation will be characterized by values of the DLE measure higher than zero. Also, the LCMV solutions for both examples are not sparse enough but reduce considerably the numbers of *False Positive* with respect to eLORETA which leads to a more precise estimation, accounting for the reprted values of the quality measures. The best results are achieved by the BC-VARETA with very sparse solution that minimizes *False Positive* estimation when compared with the other methods, in configurations when the centroids of the patches are close in distance. Also, BC-VARETA is outperforming according to the DLE values (lower values better estimation), meanwhile eLORETA and LCMV have higher localization error due to the quite smeared distribution of the solution.

**Table 1.**
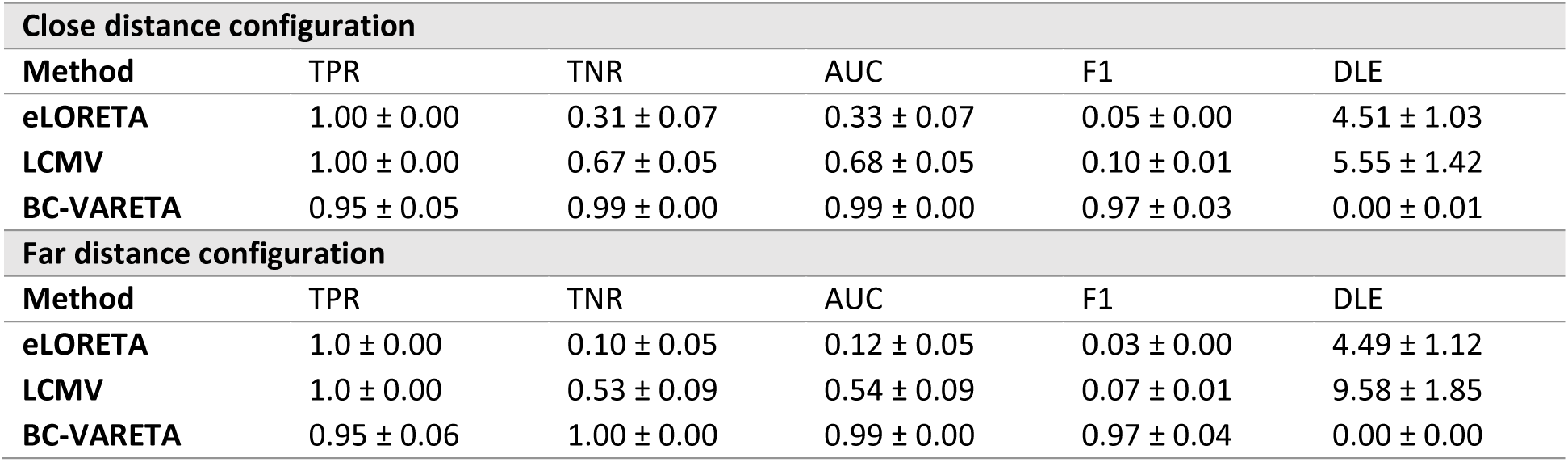
Results of ROC analysis and DLE estimation for localization performance evaluation of eLORETA, LCMV and BC-VARETA, and with the two criteria for the distance between patches: close distance (top) and far distance (bottom).

| Close distance configuration |  |  |  |  |  |
| --- | --- | --- | --- | --- | --- |
| Method | TPR | TNR | AUC | F1 | DLE |
| eLORETA | 1.00 ± 0.00 | 0.31 ± 0.07 | 0.33 ± 0.07 | 0.05 ± 0.00 | 4.51 ± 1.03 |
| LCMV | 1.00 ± 0.00 | 0.67 ± 0.05 | 0.68 ± 0.05 | 0.10 ± 0.01 | 5.55 ± 1.42 |
| BC-VARETA | 0.95 ± 0.05 | 0.99 ± 0.00 | 0.99 ± 0.00 | 0.97 ± 0.03 | 0.00 ± 0.01 |
| Far distance configuration |  |  |  |  |  |
| Method | TPR | TNR | AUC | F1 | DLE |
| eLORETA | 1.0 ± 0.00 | 0.10 ± 0.05 | 0.12 ± 0.05 | 0.03 ± 0.00 | 4.49 ± 1.12 |
| LCMV | 1.0 ± 0.00 | 0.53 ± 0.09 | 0.54 ± 0.09 | 0.07 ± 0.01 | 9.58 ± 1.85 |
| BC-VARETA | 0.95 ± 0.06 | 1.00 ± 0.00 | 0.99 ± 0.00 | 0.97 ± 0.04 | 0.00 ± 0.00 |

**Table 2.**
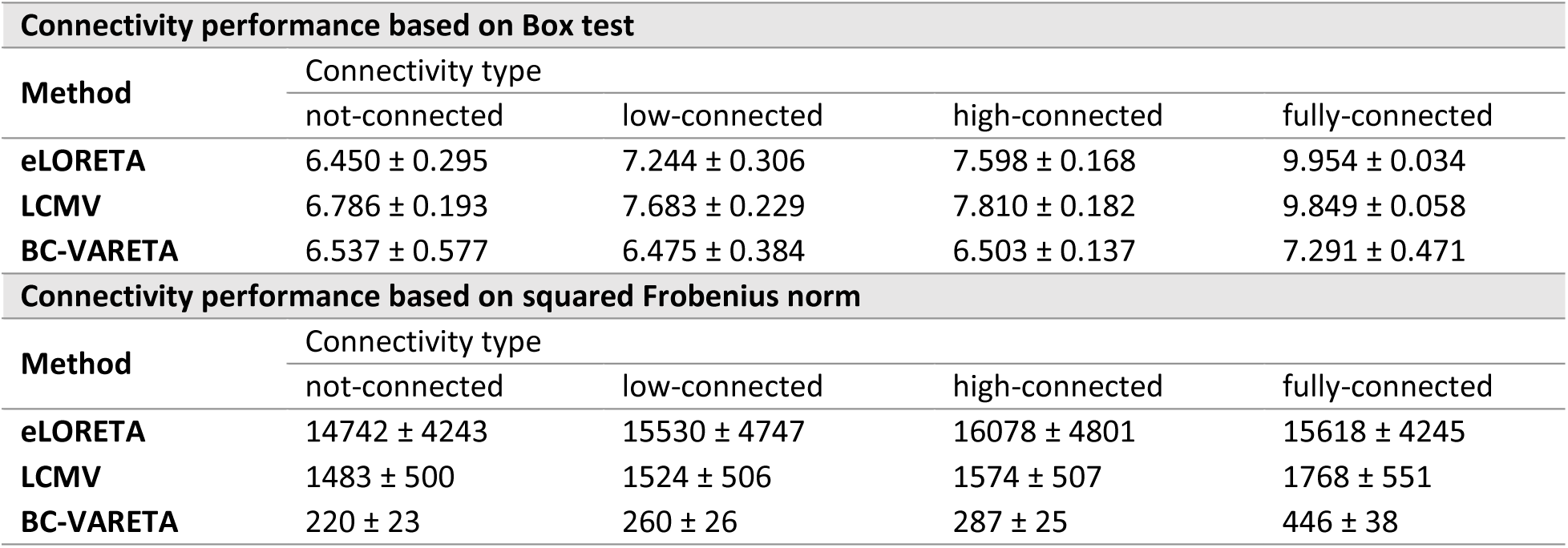
Box’s M test statistic and squared Frobenius norm for the covariance matrix estimation. The lower values represent close similitude between the simulated and estimated covariance matrices.

#### 3.2.2 Connectivity estimation performance

To evaluate BC-VARETA method performance in functional connectivity estimation, four connection modes (*not-connected*, *low-connected*, *high-connected* and *fully-connected*) were simulated based on the structure of the sources empirical covariance matrix. One example of each configuration is shown in Figure 7 with the estimation of eLORETA, LCMV and BC-VARETA. A homogeneity test between the estimated Precision Matrix (Inverse Covariance Matrix) and the simulated covariance matrix was applied, the Box’s M test statistic to evaluate the similarity between covariance matrices (*H_o_*: Σ_0_ = Σ_1_) (*Pituch and Stevens, 1994*). To contrast our results a similarity tests based on the Frobenius norm, 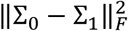 was also implemented (*TT Cai, 2017*).

**Figure 7.**
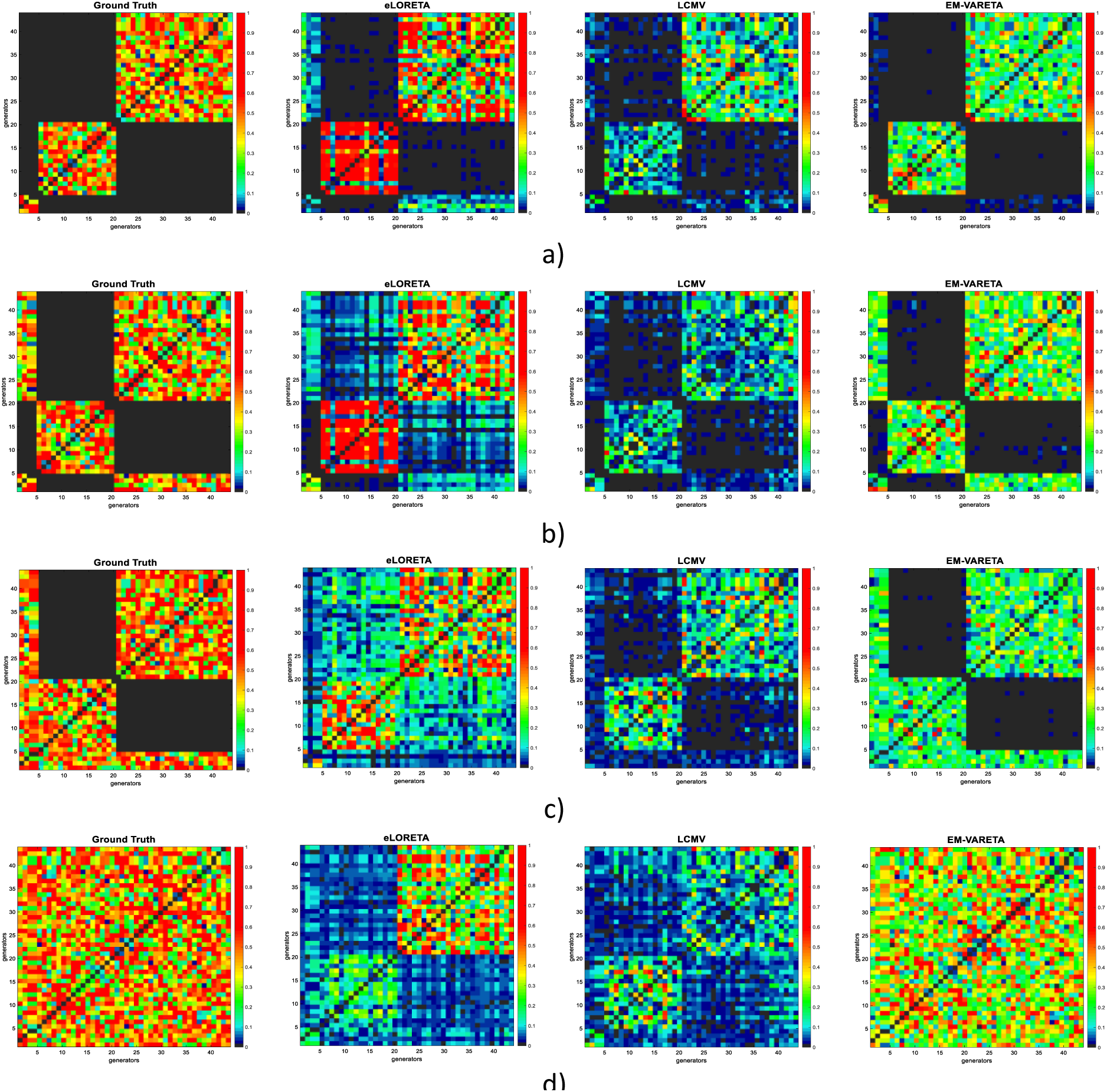
Examples of eLORETA, LCMV and FC-VARETA solution for connectivity estimation between the three patches with different size are shown. From the left to right figures represent the simulated and estimated connectivity by eLORETA, LCMV and FC-VARETA in different connectivity modes: a) not-connected, b) low-connected c) high-connected, and d) fully-connected. All the solutions were normalized and the values under a threshold of 0.01 (1%) were set to zero.

The performance of BC-VARETA over different level of SNR was evaluated and same conditions were applied to eLORETA and LCMV models to compare the results. The Figure 7 shows an example for three patches with different size under the two criteria: *far-distance* and *not-connected*.

##### Noise performance

From the results it can be noticed that with the reduction of SNR level the quality of the localization estimation as well as connectivity estimation get worse for the three methods under analysis. However, the BC-VARETA outperform over the rest in all scenarios. Furthermore, the results for 200 random patches configuration, each of them replicate at the four-connectivity mode (*not-connected*, *low-connected*, *high-connected* and *fully-connected*) and different SNR level (19 dB, 7 dB and 0 dB) are summarized in Table 3.

**Table 3.** Performance evaluation of eLORETA, LCMV and BC-VARETA based on ROC analysis (higher values represent better estimation) and DLE (lower values represent better estimation) for localization performance and the corresponding Box’s M test statistic and squared Frobenius norm of covariance matrices (lower values represent better estimation) for connectivity estimation at the three different level of noise (SNR = 19 dB, SNR = 7 dB and SNR = 0 dB).

| Localization performance based on ROC analysis and DLE |  |  |  |  |  |  |
| --- | --- | --- | --- | --- | --- | --- |
| SNR | Method | TPR | TNR | AUC | F1 | DLE |
| 19 dB | eLORETA | 1.00 ± 0.00 | 0.00 ± 0.00 | 0.02 ± 0.00 | 0.02 ± 0.00 | 3.92 ± 0.82 |
|  | LCMV | 0.98 ± 0.01 | 0.36 ± 0.04 | 0.37 ± 0.04 | 0.06 ± 0.00 | 22.81 ± 2.41 |
|  | BC-VARETA | 0.84 ± 0.06 | 0.93 ± 0.01 | 0.92 ± 0.01 | 0.36 ± 0.06 | 5.28 ± 1.67 |
| 7 dB | eLORETA | 1.00 ± 0.00 | 0.00 ± 0.00 | 0.02 ± 0.00 | 0.04 ± 0.00 | 4.15 ± 1.00 |
|  | LCMV | 0.85 ± 0.07 | 0.32 ± 0.04 | 0.33 ± 0.04 | 0.05 ± 0.04 | 25.42 ± 2.46 |
|  | BC-VARETA | 0.74 ± 0.08 | 0.92 ± 0.01 | 0.91 ± 0.01 | 0.28 ± 0.04 | 7.78 ± 2.26 |
| 0 dB | eLORETA | 1.00 ± 0.00 | 0.00 ± 0.00 | 0.01 ± 0.00 | 0.03 ± 0.00 | 12.65 ± 2.91 |
|  | LCMV | 0.72 ± 0.12 | 0.29 ± 0.04 | 0.30 ± 0.04 | 0.03 ± 0.00 | 30.65 ± 3.35 |
|  | BC-VARETA | 0.65 ± 0.11 | 0.92 ± 0.01 | 0.91 ± 0.01 | 0.22 ± 0.04 | 9.98 ± 2.59 |
| Connectivity performance based on Box's M test statistic |  |  |  |  |  |  |
| SNR | Method | Connectivity type |  |  |  |  |
|  |  | not connected | low connected | high connected | fully connected |  |
| 19 dB | eLORETA | 6.471 ± 0.304 | 7.216 ± 0.287 | 7.621 ± 0.224 | 9.950 ± 0.037 |  |
|  | LCMV | 6.506 ± 0.173 | 6.820 ± 0.178 | 7.660 ± 0.146 | 9.883 ± 0.046 |  |
|  | BC-VARETA | 6.105 ± 0.676 | 6.443 ± 0.575 | 6.784 ± 0.550 | 7.473 ± 0.953 |  |
| 7 dB | eLORETA | 6.367 ± 0.247 | 7.197 ± 0.249 | 7.630 ± 0.218 | 9.923 ± 0.050 |  |
|  | LCMV | 6.240 ± 0.107 | 6.239 ± 0.129 | 7.541 ± 0.112 | 9.899 ± 0.045 |  |
|  | BC-VARETA | 5.117 ± 0.806 | 5.608 ± 0.726 | 5.750 ± 0.698 | 6.356 ± 1.176 |  |
| 0 dB | eLORETA | 6.222 ± 0.149 | 7.117 ± 0.150 | 7.564 ± 0.136 | 9.865 ± 0.071 |  |
|  | LCMV | 6.083 ± 0.069 | 5.780 ± 0.075 | 7.561 ± 0.080 | 9.909 ± 0.042 |  |
|  | BC-VARETA | 4.488 ± 1.075 | 4.694 ± 1.107 | 4.783 ± 0.955 | 4.682 ± 1.249 |  |
| Connectivity performance based on Frobenius norm |  |  |  |  |  |  |
| SNR | Method | Connectivity type |  |  |  |  |
|  |  | not connected | low connected | high connected | fully connected |  |
| 19 dB | eLORETA | 15843 ± 4127 | 16047 ± 4603 | 16276 ± 4545 | 16241 ± 4153 |  |
|  | LCMV | 933 ± 97 | 974 ± 96 | 984 ± 96 | 1151 ± 105 |  |
|  | BC-VARETA | 212 ± 20 | 254 ± 24 | 279 ± 24 | 440 ± 33 |  |
| 7 dB | eLORETA | 19949 ± 4682 | 21372 ± 5692 | 21065 ± 5128 | 21522 ± 5580 |  |
|  | LCMV | 843 ± 69 | 882 ± 78 | 916 ± 78 | 1060 ± 78 |  |
|  | BC-VARETA | 210 ± 20 | 252 ± 25 | 277 ± 25 | 437 ± 33 |  |
| 0 dB | eLORETA | 31375 ± 5691 | 30763 ± 5179 | 31291 ± 5453 | 31091 ± 5082 |  |
|  | LCMV | 834 ± 66 | 863 ± 71 | 902 ± 63 | 1053 ± 74 |  |
|  | BC-VARETA | 213 ± 32 | 250 ± 30 | 278 ± 30 | 437 ± 38 |  |

### 3.3 Real data analysis

#### 3.3.1 Real EEG data

The real EEG data was gathered as part of the Cuban Human Brain Mapping Project (*Hernandez-Gonzalez, 2011*) created in 2005 with the aim to obtain atlases for normal and pathological Cuban population. The EEG data case under analysis in this research belong to healthy male subject of 32 years old in resting state condition with eyes closed. The EEG was recorded using a MEDICID 5 EEG recording system of 128 channels with sampling frequency of 200 Hz.

The physiology of human EEG signal for resting condition, with eyes-closed and eyes-open, have been wide studied and its spectral characteristic and brain areas associated are well-defined (*Barry, 2007*): delta band is related to frontal activation and alfa band is related to occipital activation. In Figure 8 the estimation of BC-VARETA for two specific frequency bins, 1.57 Hz and 10.57 Hz, that belongs to delta (1.5 Hz – 3.5 Hz) and alpha (8 Hz – 13 Hz), respectively. Here the BC-VARETA results shown a high corresponding with the state of art studies on the physiology of difference frequency band of the EEG spectra.

**Figure 8.**
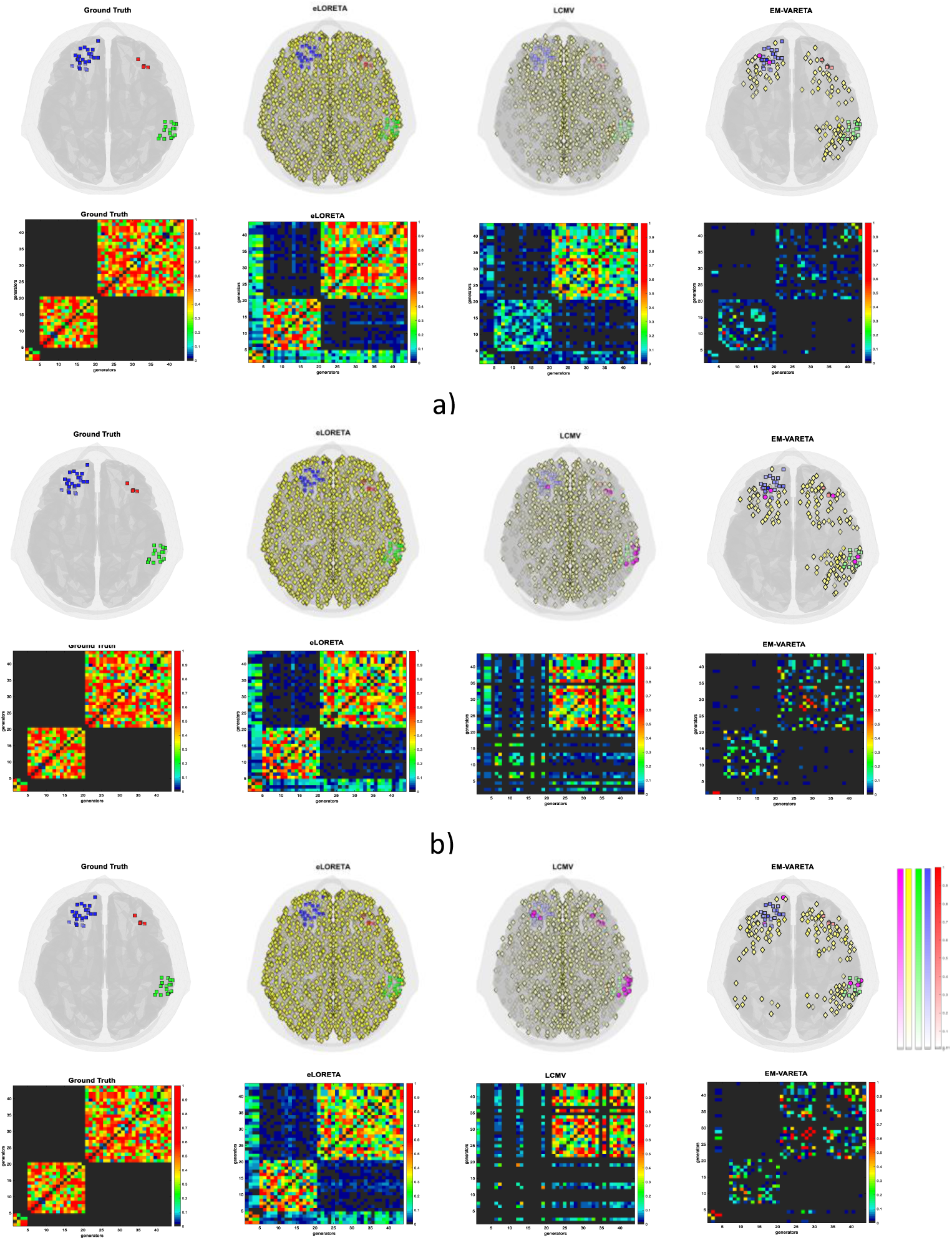
Three none-connected patches with different size and the corresponding eLORETA, LCMV and FC-VARETA estimation. Three level of noise were applied: a) SNR = 19 dB, b) SNR = 7 dB and c) SNR = 0 dB. The small squares drew at cortical surface represents the True Positive estimation of the three patches differentiated by colors (red color for patch 1, green color for patch 2 and blue color for patch 3), the small yellow diamonds represent the False Positive estimation and the magenta circles represent the False Negative estimation. All the solutions were normalized and the values under a threshold of 0.01 (1%) were set to zero.

**Figure 9.**
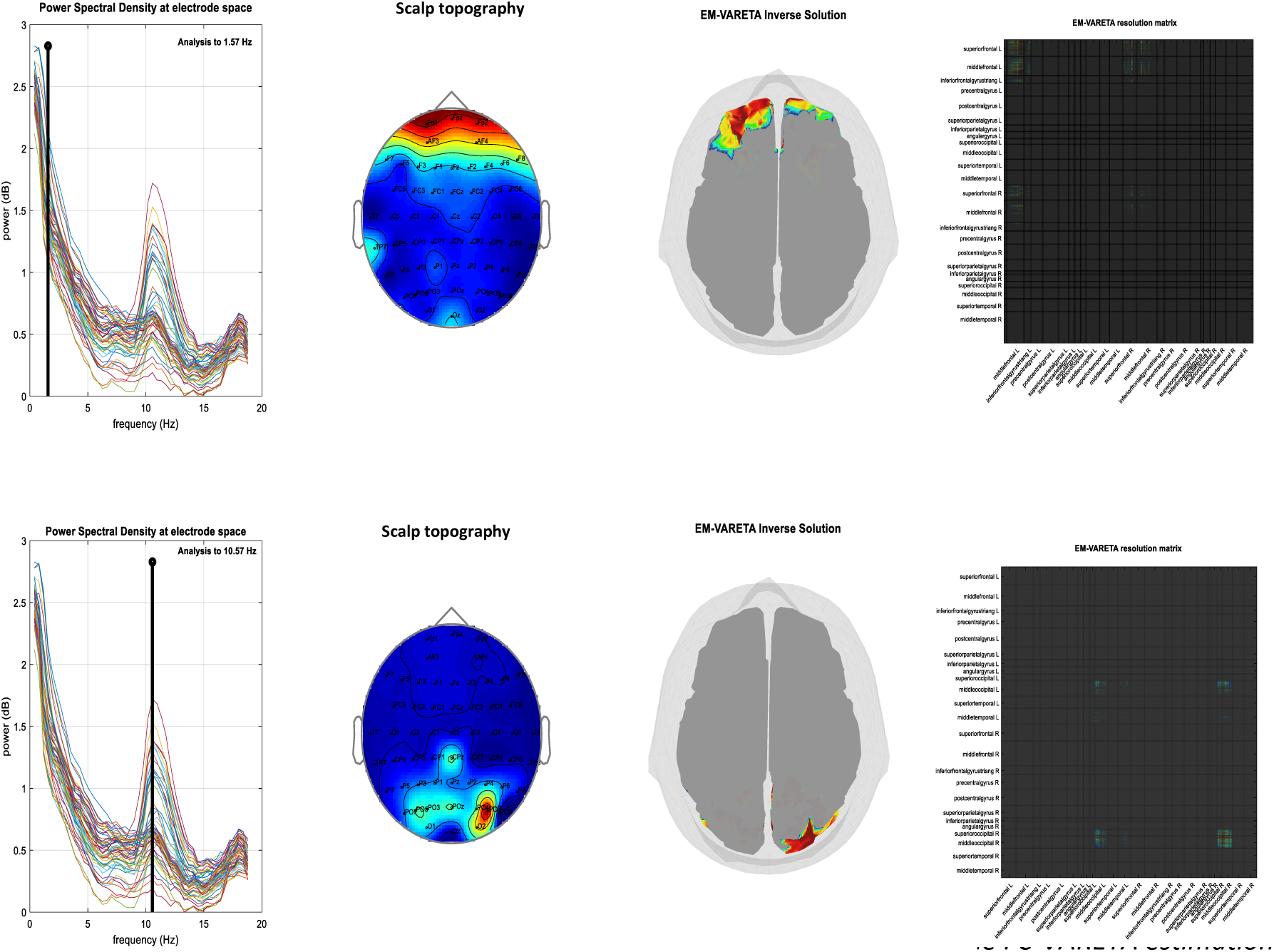
Real EEG data for a healthy subject with eyes closed in resting state condition. The FC-VARETA estimation was performed at two frequency bins: top) 1.57 Hz and bottom) 10.57 Hz, which represent the delta and alpha bands respectively. From left to right is showed the power spectral density for each of 58 EEG channels, the scalp topography of 58 channels for a specific bin of frequency, the localization estimation performed by BC-VARETA and the connectivity estimation between anatomical areas of the brain.

#### 3.3.2 Real MEG data

MEG real data of a healthy adult was picked from the WU-Minn HCP consortium in the MEG Initial Data Release, the data corresponds to the resting condition with eyes open and attention fixation on a projected red crosshair (*Marcus et.al., 2011*). This data was recorded in 248 magnetometer channels of a MEG system for three runs of approximately 5 minutes each, at a sampling rate of 508 Hz. Analogously the previous EEG study, the BC-VARETA activity and connectivity analysis was performed at the delta and alpha bands. The results are shown in Figure 10.

**Figure 10.**
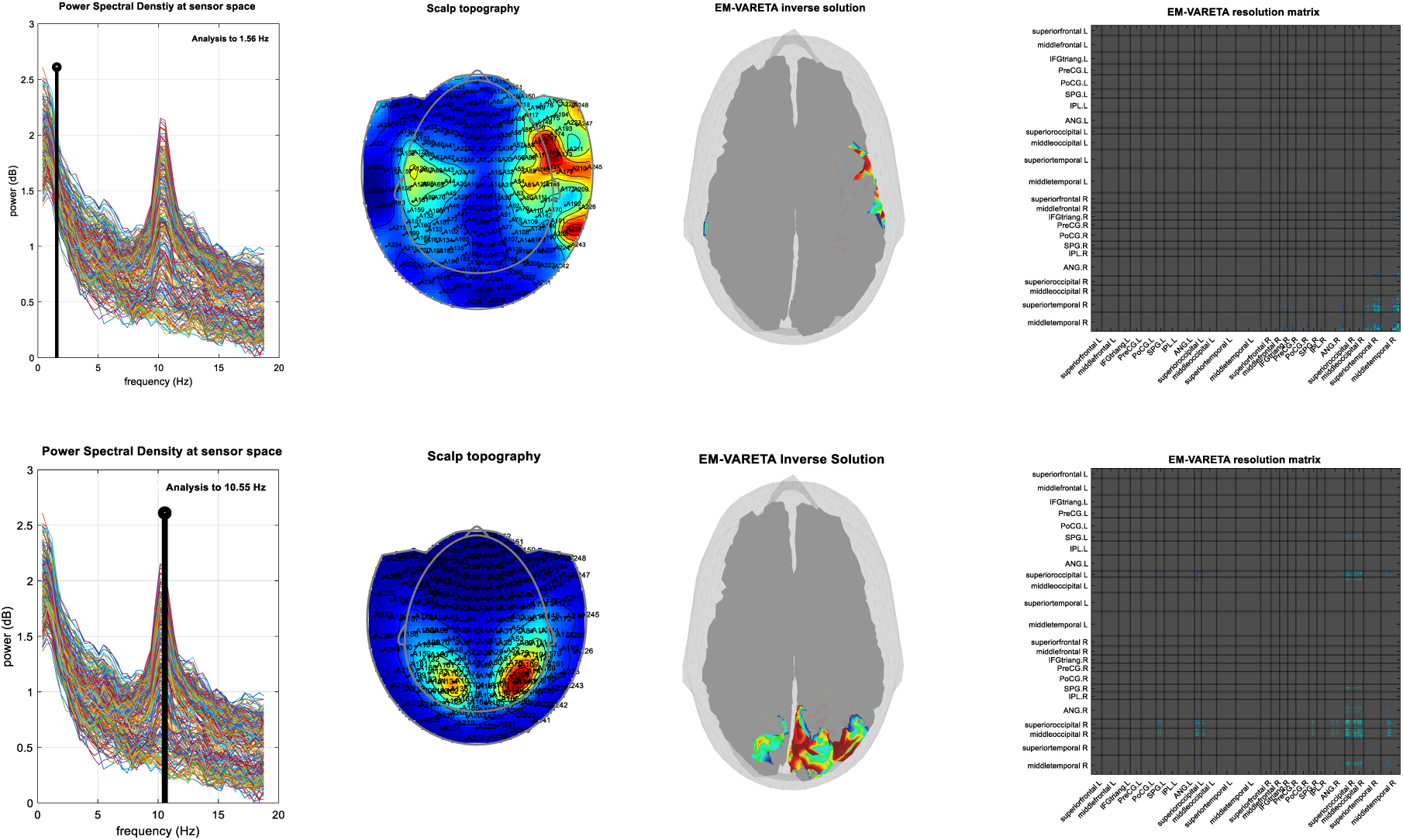
Real MEG data for a healthy subject with open eyes in resting state condition. The FC-VARETA estimation was performed at two frequency bins: top) 1.55 Hz and bottom) 10.55 Hz, which represent the delta and alpha bands respectively. From left to right it is shown the power spectral density for each of 248 MEG sensors, the scalp topography of 248 sensors for an specific frequency bin, the localization estimation by BC-VARETA and the connectivity estimation between anatomical areas of the brain.

## 4 CONCLUSIONS

A novel methodology, BC-VARETA, for JOINT ESTIMATION of Sources ACTIVITY and its CONNECTIVITY parameters is described. SUPER-RESOLUTION in the connectivity estimation through Sparse Hermitian Sources Graphical Model is achieved.

## 5 ACKNOWLEDGEMENT

This study was funded by the Grant No. 61673090 from the National Nature Science Foundation of China.

## Footnotes

1 The superscript ^T^ will denote transpose and the superscript -1 will denotes matrix inversion across the document.

2 Here the symbol ⊙ represents the element wise matrix product (Hadamard) and sign(**Θ*_JJ_***) is the element wise

